# Sex differences in sensitivity to dopamine receptor manipulations of risk-based decision making in rats

**DOI:** 10.1101/2024.04.18.590124

**Authors:** Alexa-Rae Wheeler, Leah M. Truckenbrod, Adrian Boehnke, Caitlin A. Orsini

## Abstract

Risky decision making involves the ability to weigh risks and rewards associated with different options to make adaptive choices. Previous work has established a necessary role for the basolateral amygdala (BLA) in mediating effective decision making under risk of punishment, but the mechanisms by which the BLA mediates this process are less clear. Because this form of decision making is profoundly sensitive to dopaminergic (DA) manipulations, we hypothesized that DA receptors in the BLA may be involved in risk-taking behavior. To test this hypothesis, male and female Long-Evans rats were trained in a decision-making task in which rats chose between a small, safe food reward and a larger food reward that was associated with a variable risk of footshock punishment. Once behavioral stability emerged, rats received intra-BLA infusions of ligands targeting distinct dopamine receptor subtypes prior to behavioral testing. Intra-BLA infusions of the dopamine D2 receptor (D2R) agonist quinpirole decreased risk taking in females at all doses, and this reduction in risk taking was accompanied by an increase in sensitivity to punishment. In males, decreased risk taking was only observed at the highest dose of quinpirole. In contrast, intra-BLA manipulations of dopamine D1 or D3 receptors (D1R and D3R, respectively) had no effect on risk taking. Considered together, these data suggest that differential D2R sensitivity in the BLA may contribute to the well-established sex differences in risk taking. Neither D1Rs nor D3Rs, however, appear to contribute to risky decision making in either sex.

## Introduction

The basolateral amygdala (BLA) is a critical component of the neural network that governs risk-based decision making. Indeed, BLA damage in humans results in greater risk-taking behavior in laboratory gambling tasks designed to simulate real-world risk-based decision making [1,2]. Neuroimaging studies have reported greater activity in the amygdala following choices associated with potential losses [3] as well as greater gray matter volume in individuals who display higher risk tolerance [4]. Recent work using rodent models of decision making have also used a range of approaches to systematically dissect the contribution of the BLA to different forms of risk-based decision making [5–10]. For example, pharmacological inactivation of the BLA decreases risky choice in a decision-making task in which the large reward is associated with risk of reward omission [9]. In contrast, permanent lesions of the BLA increase risky choice in a decision-making task that involves risk of explicit punishment [8]. Optogenetic approaches have further refined our understanding of the role of the BLA in this latter form of decision making by showing that BLA activity contributes to decision making under risk of punishment differently across the decision process [11]. Collectively, these studies have established that the BLA is necessary for the integration of reward- and risk-related information to guide adaptive choice behavior. Very little is known, however, about the distinct receptor mechanisms within the BLA that are involved in risk-based decision making, particularly when the risk is that of potential punishment.

Decision making involving risk of explicit punishment is exquisitely sensitive to dopaminergic (DA) manipulations [12–14]. For instance, systemic administration of the indirect DA agonist amphetamine [12,14,15] or DA D2 receptor (D2R) agonists decrease risky choice [12,13]. Microinfusions of D2R agonists directly into the nucleus accumbens similarly decrease risky choice [16]. Despite evidence that the BLA receives DA input from midbrain regions [17–19] and that DA receptors (and D2Rs in particular) in the BLA are important for dopamine’s ability to gate BLA activity [20,21], very few studies have in fact investigated the role of BLA DA receptors in decision making under risk of punishment. This is surprising in light of the fact that both aversive and appetitive stimuli elicit DA release in the BLA [22–26] and that DA receptor manipulations in the BLA impact reward- and avoidance-related behavior [27,28]. Moreover, using a probability discounting task, Larkin et al. (2016) reported that both dopamine D1 receptors (D1R) and D2Rs in the BLA are involved in decision making involving reward uncertainty. Although the neurobiological substrates of decision making can differ as a function of the risk involved [30,31], these data provide further support for the role of BLA DA receptors in decision making under risk of punishment.

The objective of the current study was to determine the contribution of DA receptors in the BLA to decision making under risk of explicit punishment. To model this form of decision making, rats choose between a small, safe food reward and a large food reward that is accompanied by potential footshock delivery, the probability of which systematically increases across the session. Once rats were trained in this task, they received intra-BLA infusions of DA receptor ligands prior to behavioral testing. The findings from these experiments reveal novel information about the mechanisms by which DA can modulate BLA activity in a sex-dependent manner.

## Materials and Methods

### Subjects

Male (n=37) and female (n=38) Long-Evans rats were housed individually and maintained on a 12-hour reverse light/dark cycle (lights off at 0800). Rats were given free access to food and water during their acclimation period. During behavioral testing, however, rats were food restricted to 85% of their free-feeding weight. Water was provided *ad libitum*. All procedures were conducted in accordance with The University of Texas at Austin Institutional Animal Care and Use Committee and adhered to the ethical guidelines of the National Institutes of Health.

### Overview of experimental design

In all experiments, rats first underwent surgery to implant bilateral cannulae targeting the BLA and then were trained on the Risky Decision-making Task (RDT).

The goal of Experiment 1 was to determine the role of D2Rs in the BLA in decision making involving risk of explicit punishment. In Experiment 1.1, male and female rats were trained on the RDT until stable choice performance was achieved, after which the effects of intra-BLA microinfusions of the D2/3 receptor agonist quinpirole were evaluated. Guided by the results from Experiment 1.1., Experiment 1.2 was conducted using an identical experimental design as Experiment 1.1, but a higher dose of quinpirole was infused into the BLA. In Experiment 1.3, the experimental procedures were the same as those in Experiments 1.1 and 1.2, except rats received intra-BLA microinfusions of the D2R antagonist eticlopride.

Because quinpirole binds to both D2Rs and D3 dopamine receptors (D3Rs), Experiment 2 was conducted to ascertain whether effects of quinpirole on risk taking in Experiments 1.1 and 1.2 were due to D2R or D3R activation. After reaching behavioral stability, rats received intra-BLA microinfusions of the selective D3R agonist PD128907 prior to being tested on the RDT.

The goal of Experiment 3 was to identify the role of D1Rs in the BLA in decision making involving risk of explicit punishment. In Experiment 3.1, male and female rats were trained on the RDT until they achieved behavioral stability, after which the effects of intra-BLA infusions of the D1R agonist SKF81297 on risk taking were examined. The experimental design of Experiment 3.2 was identical to that of Experiment 3.1, except rats received intra-BLA microinfusions of the D1R antagonist SCH23390.

### Surgical procedures

Details of the surgical procedures are found in the Supplemental Information. Briefly, rats underwent stereotaxic surgery in which bilateral cannulae were implanted directly above the BLA. Rats were given one week to recover from surgery, after which they were food restricted in preparation for behavioral training.

### Risky Decision-making Task

Behavioral training was conducted in 6 operant chambers, each of which was outfitted with 2 retractable levers flanking a centrally located food trough. A nosepoke hole was located directly above the food trough. The floor consisted of stainless-steel rods through which scrambled shocks were delivered. Rats were initially trained to perform individual components of the decision-making task, such as nosepoking and pressing levers for food. Upon completion of behavioral shaping, rats progressed to the decision-making task in which they were presented with either one lever (forced choice trials) or two levers (free choice trials). A press on one lever yielded a small, safe food reward (1 pellet) whereas a press on the other lever yielded a large food reward (2 pellets) accompanied by a risk of mild footshock punishment (1 s). The probability of footshock delivery was specific to each of the 5 trial blocks and systematically increased across the 60-min session (0, 25, 50, 75, 100%).

### Microinfusions

Rats received intra-BLA microinjections (0.5 µl at a rate of 0.4 µl/min in each hemisphere) of quinpirole (D2/3R agonist; Experiments 1.1 and 1.2), eticlopride (D2R antagonist; Experiment 1.3), PD128907 (D3R agonist; Experiment 2), SKF81297 (D2R agonist; Experiment 3.1), or SCH23390 (D1R antagonist; Experiment 3.2). In each experiment, the different doses of each drug were microinfused using a randomized, within-subjects Latin-square design such that each rat received each dose of the drug and its corresponding vehicle, with a minimum of 48 hours between each successive round of microinfusions.

### Data analysis

Detailed descriptions of the statistical analyses used to analyze the behavioral data are provided in the Supplemental Information. The primary dependent variable in the RDT was the choice of the large, risky lever (i.e., risk taking). Effects of the selective DA receptor ligands on risk taking were assessed using a repeated-measures ANOVA (RMANOVA), with trial block and drug dose as within-subjects factors and sex as the between-subjects factor. Results were deemed significant when *p* ≤ 0.05. If parent ANOVAs yielded main effects (dose, sex) or significant interactions (dose X trial block, dose X sex, dose X trial block X sex), additional RMANOVAs were conducted to determine the source of the significance. In these instances, Bonferroni’s corrections were used to adjust *p*-values to account for multiple comparisons.

## Results

### Experiment 1: The role of D2Rs in the BLA in risk taking

#### Experiment 1.1

A three-factor RMANOVA revealed that quinpirole dose-dependently decreased overall choice of the large, risky reward [dose, *F*(2,34)=12.14, *p*<0.01; ƞ^2^=0.42; dose X trial block, *F*(8,136)=0.49, *p*=0.86, ƞ^2^=0.03]. Although there was no dose X sex X trial block interaction [*F*(8,136)=0.87, *p*=0.54, ƞ^2^=0.05], there was a significant interaction between dose and sex [*F*(2,24)=6.94, *p*<0.01, ƞ^2^=0.29], suggesting that quinpirole affected choice of the large risky reward, irrespective of trial block, in a sex-dependent manner. As shown in Figure 1, this interaction appeared to be driven by an effect of quinpirole in females, but not males. Indeed, when performance in the RDT was compared between doses in males and females separately, quinpirole dose-dependently decreased choice of the large, risky reward in females [dose, *F*(2,16)=15.62, *p*<0.01, ƞ^2^=0.66; dose X trial block, *F*(8, 64)=0.65, *p*=0.73, ƞ^2^=0.08] but had no effect in males [dose, *F*(2,18)=0.95, *p=*0.41, ƞ^2^=0.10; dose X trial block, *F*(8,72)=0.66, *p*=0.72, ƞ^2^=0.07]. Post-hoc comparisons in females showed that both the low [dose, *F*(1,8)=17.18, *p*<0.01, ƞ^2^=0.68; dose X trial block, *F*(4,32)=0.88, *p*=0.49, ƞ^2^=0.10] and the high [dose, *F*(1, 9)=20.70, *p*<0.01, ƞ^2^=0.70; dose X trial block, *F*(4,36)=0.42, *p*=0.80, ƞ^2^=0.04] dose of quinpirole reduced choice of the large, risky reward relative to vehicle. Given that quinpirole altered performance only in females, subsequent analyses of win-stay and lose-shift performance were restricted to this sex. Although there was no main effect of dose [*F*(2,12)=1.15, *p*=0.35, ƞ^2^=0.16], there was a significant dose X trial type (win-stay vs. lose-shift) interaction [*F*(2,12)=5.98, *p*=0.02, ƞ^2^=0.50]. To determine the source of this significance, win-stay and lose-shift were separately compared across doses. Although Figure 1C suggests that both the low and high dose of quinpirole decreased win-stay behavior, this analysis did not survive correction for multiple comparisons [*F*(2,18)=3.67, *p*=0.05, ƞ^2^=0.29]. In contrast, there was a main effect of dose when comparing the proportion of lose-shift trials across quinpirole doses [*F*(2,16)=20.45, *p*<0.01, ƞ^2^=0.79], with the proportion of these trials increasing at the highest dose (*p*<0.001).

**Figure 1.**
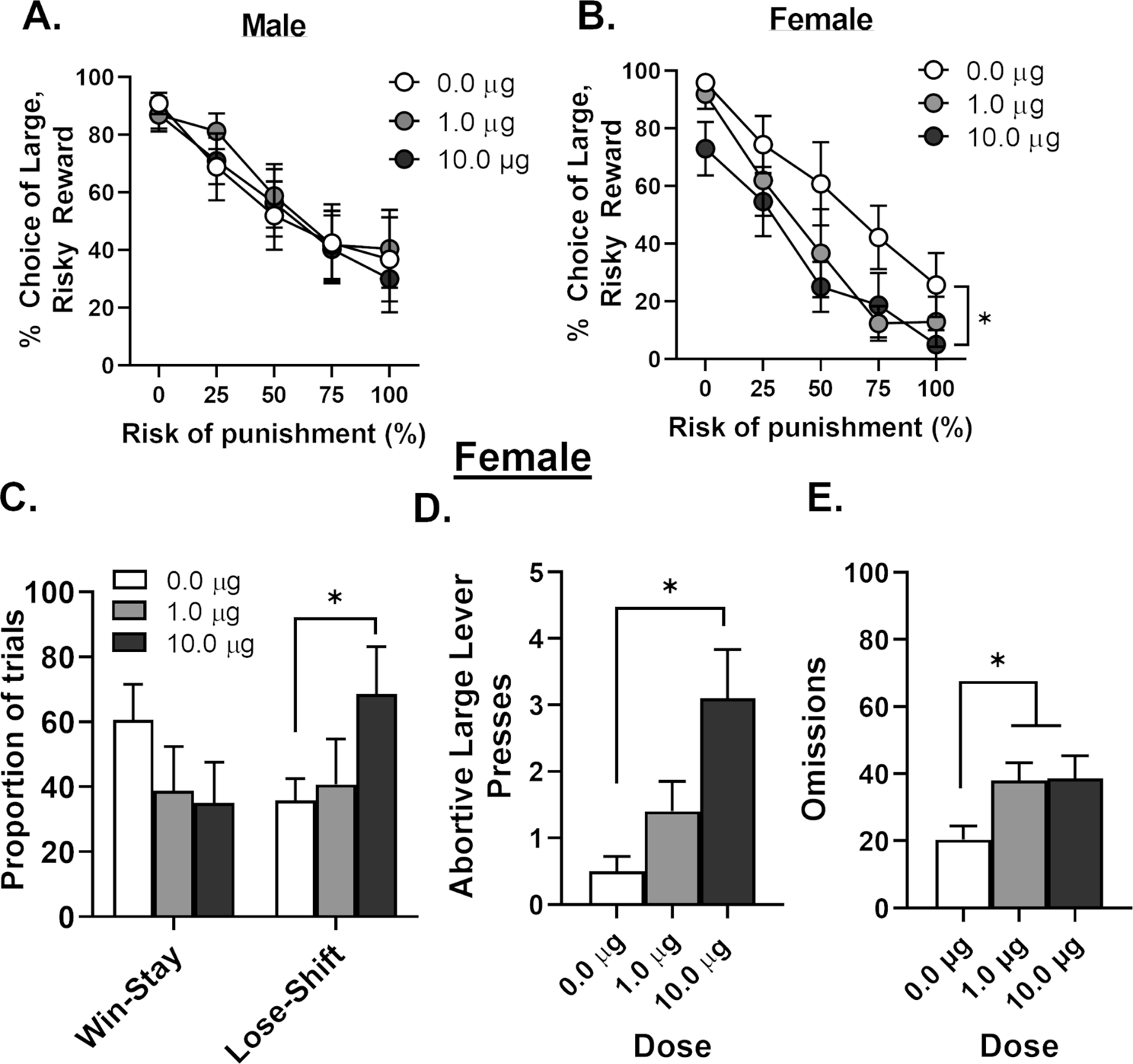
Effects of intra-BLA infusions of a D2R agonist on risk taking. **A.** In males, intra-BLA infusions of the D2R agonist quinpirole did not alter choice of the large, risky reward relative to vehicle infusions. **B.** In females, intra-BLA infusions of quinpirole dose-dependently decreased choice of the large, risky reward relative to vehicle infusions. Absence of error bars (e.g., Block 1, vehicle dose) is because the size of the standard error of the mean (SEM) was smaller than the point displayed. **C.** Whereas there was no effect of quinpirole on the proportion of win-stay trials, the highest dose of quinpirole caused a significant increase in the proportion of lose-shift trials in females. **D.** Intra-BLA infusions of the highest dose of quinpirole increase the number of abortive large lever presses in females. **E.** Intra-BLA infusions of both the low and high doses of quinpirole increased the percentage of omitted free choice trials. Data are presented as mean ± SEM. Asterisks denote statistical significance (*p* < 0.05).

During the period in which rats deliberated between the two available options, rats exhibited “abortive lever pressing” for the lever associated with the large, risky lever. Variants of this behavior have been characterized previously as one behavioral index of risk assessment [32–35]. In the RDT, abortive lever pressing was defined as physical contact with the large, risky lever (paw or mouth contact or sniffing) followed by either a press on the small, safe lever or complete omission of the trial. Because abortive lever presses were only observed in blocks 2 through 5, they were summed across these four blocks and then compared between doses in females using a one-factor RMANOVA. As evident in Figure 1D, quinpirole increased the number of abortive lever presses [*F*(2,18)=7.51, *p*<0.01, ƞ^2^=0.45]. Post-hoc comparisons between vehicle and each dose revealed that this increase was driven by the high dose of quinpirole [high: *t*(18)=3.82, *p*<0.01; low dose: *t*(18)=1.32, *p*=0.61]. Finally, analyses of the percentage of omitted free choice trials revealed that although quinpirole did not alter omissions in males [*F*(2,20)=1.68, *p*=0.21, ƞ^2^=0.14], both the low and high doses increased omissions in females [*F*(2,18)=6.52, *p*<0.01, ƞ^2^=0.42; Figure 1E].

#### Experiment 1.2

Experiment 1.1 suggests that the role of BLA D2Rs in risk taking is sex dependent. It is conceivable, however, that BLA D2Rs of females are just more sensitive to D2R agonists compared with males and that a higher dose of quinpirole is necessary to modulate risk taking in males. Consequently, male and female rats received intra-BLA infusions of a higher dose of quinpirole (40 µg) and were tested in the RDT. Quinpirole once again reduced choice of the large, risky reward [dose, *F*(1,11)=11.24, *p*<0.01, ƞ^2^=0.51; dose X trial block, *F*(4,44)=2.68, *p*=0.04, ƞ^2^=0.20], but this effect was now observed in both male and female rats [dose X sex, *F*(1,11)=0.01, *p*=0.92, ƞ^2^<0.01; dose X sex X trial block, *F*(4,44)=1.78, *p*=0.15, ƞ^2^=0.14; Figure 2A,B]. There were, however, no effects of quinpirole on win-stay or lose-shift behavior (collapsed across sex; dose, *F*(1,9)=1.49, *p*=0.25, ƞ^2^=0.14; dose X trial type, *F*(1,9)=1.62, *p*=0.24, ƞ^2^=0.15]. Although this is inconsistent with results from Experiment 1.1, it is likely due to the smaller sample size used for Experiment 1.2, and the fact that quinpirole caused several rats (particularly females) to exclusively choose the small reward, precluding the ability to categorize trials as win-stay or lose-shift in those rats. As depicted in Figure 2C, the high dose of quinpirole increased the number of abortive lever presses relative to vehicle in both male and female rats [dose, *F*(1,11)=25.49, *p*<0.01, ƞ^2^=0.70; sex, *F*(1,11)=0.25, *p*=0.62, ƞ^2^=0.02; dose X sex, *F*(1,11)=0.69, *p*=0.43, ƞ^2^=0.06]. In contrast, the higher dose of quinpirole did not alter the percentage of omitted free choice trials in males or females [dose, *F*(1,11)=3.84, *p*=0.08, ƞ^2^=0.26; sex, *F*(1,11)=3.07, *p*=0.11, ƞ^2^<0.01; dose X sex, *F*(1,11)<0.01, *p*=0.95, ƞ^2^<0.01; Figure 2D].

**Figure 2.**
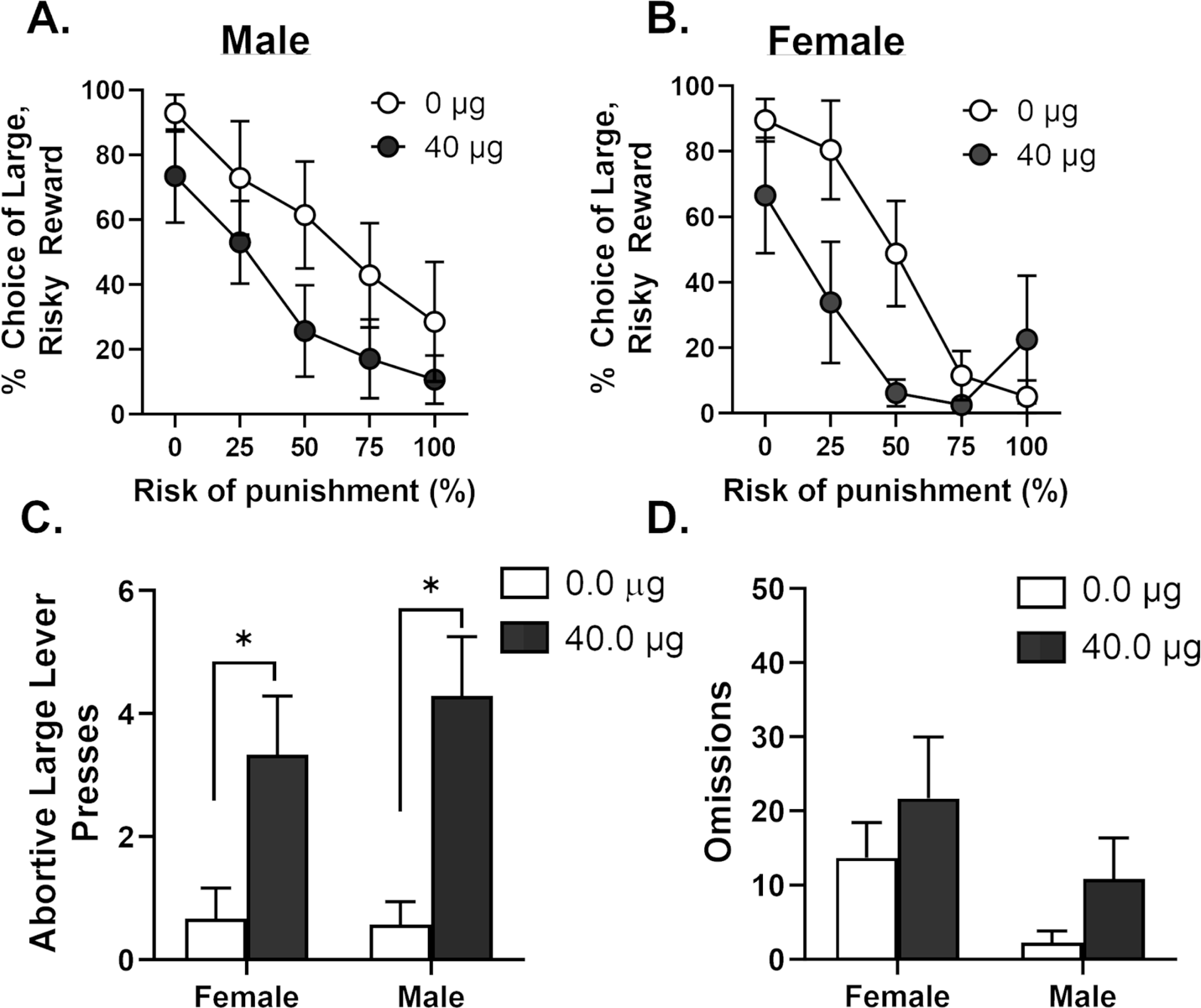
Effects of intra-BLA infusions of a higher dose of quinpirole on risk taking. **A.** In males, intra-BLA infusions of a higher dose of quinpirole (40 µg) decreased choice of the large, risky reward relative to vehicle infusions. **B.** In females, intra-BLA infusions of a higher dose of quinpirole also decreased choice of the large, risky reward relative to vehicle infusions. **C.** The higher dose of quinpirole increased the number of abortive large lever presses in both males and females. **D.** Intra-BLA infusions of the higher dose of quinpirole did not reliably affect the percentage of omitted free choice trials in males or females. Data are presented as mean ± standard error of the mean. Asterisks denote statistical significance (*p* < 0.05).

#### Experiment 1.3

Intra-BLA microinjections of eticlopride did not affect choice of the large, risky reward [dose, *F*(2,32)=0.20, *p*=0.82, ƞ^2^=0.01; dose X trial block, *F*(8,128)=1.40, *p*=0.20, ƞ^2^=0.08] in males or females [sex, *F*(1,16)=0.33, *p*=0.57, ƞ^2^=0.02; dose X sex, *F*(2,32)=0.34, *p*=0.72, ƞ^2^=0.02; dose X sex X trial block, *F*(8,128)=0.76, *p*=0.64; ƞ^2^=0.05; Figure 3].

**Figure 3.**
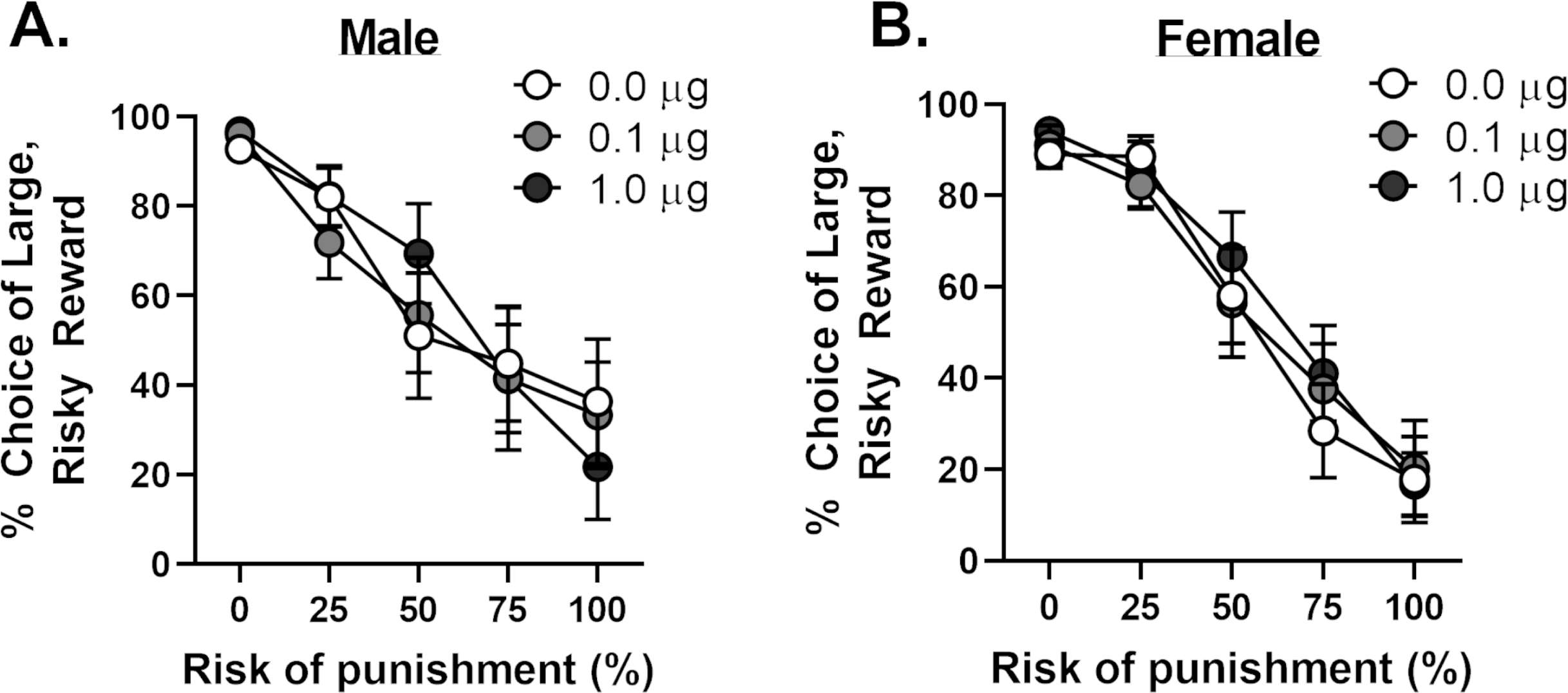
Effects of intra-BLA infusions of a D2R antagonist on risk taking. **A.** In males, intra-BLA infusions of the D2R antagonist eticlopride did not alter choice of the large, risky reward relative to vehicle infusions. **B.** In females, intra-BLA infusions of eticlopride did not alter choice of the large, risky reward relative to vehicle infusions. Data are presented as mean ± standard error of the mean (SEM). Absence of error bars (e.g., Block 1) is because the size of the SEM was smaller than the point displayed.

### Experiment 2: The role of BLA D3Rs in risk taking

Because quinpirole can bind to both D2Rs and D3Rs, it is conceivable that its ability to reduce risk taking in females was due to activity at D3Rs. To address this possibility, rats received intra-BLA infusions of the selective D3R agonist PD128907 and were tested in the RDT. In contrast to the effects of quinpirole, PD128907 did not affect performance in the RDT [dose, *F*(2,32)=0.66, *p*=0.52, ƞ^2^=0.04; dose X trial block, *F*(8,128)=1.28, *p*=0.26, ƞ^2^=0.07] in males or females [sex, *F*(1,16)=2.13, *p*=0.16; ƞ^2^=0.12; dose X sex, *F*(2,32)=0.07, *p*=0.93, ƞ^2^<0.01; dose X sex X trial block, *F*(8,128)=1.28, *p*=0.26, ƞ^2^=0.07; Figure 4]. These results suggest that quinpirole likely reduced risk taking through activation of D2Rs, but not D3Rs, in the BLA.

**Figure 4.**
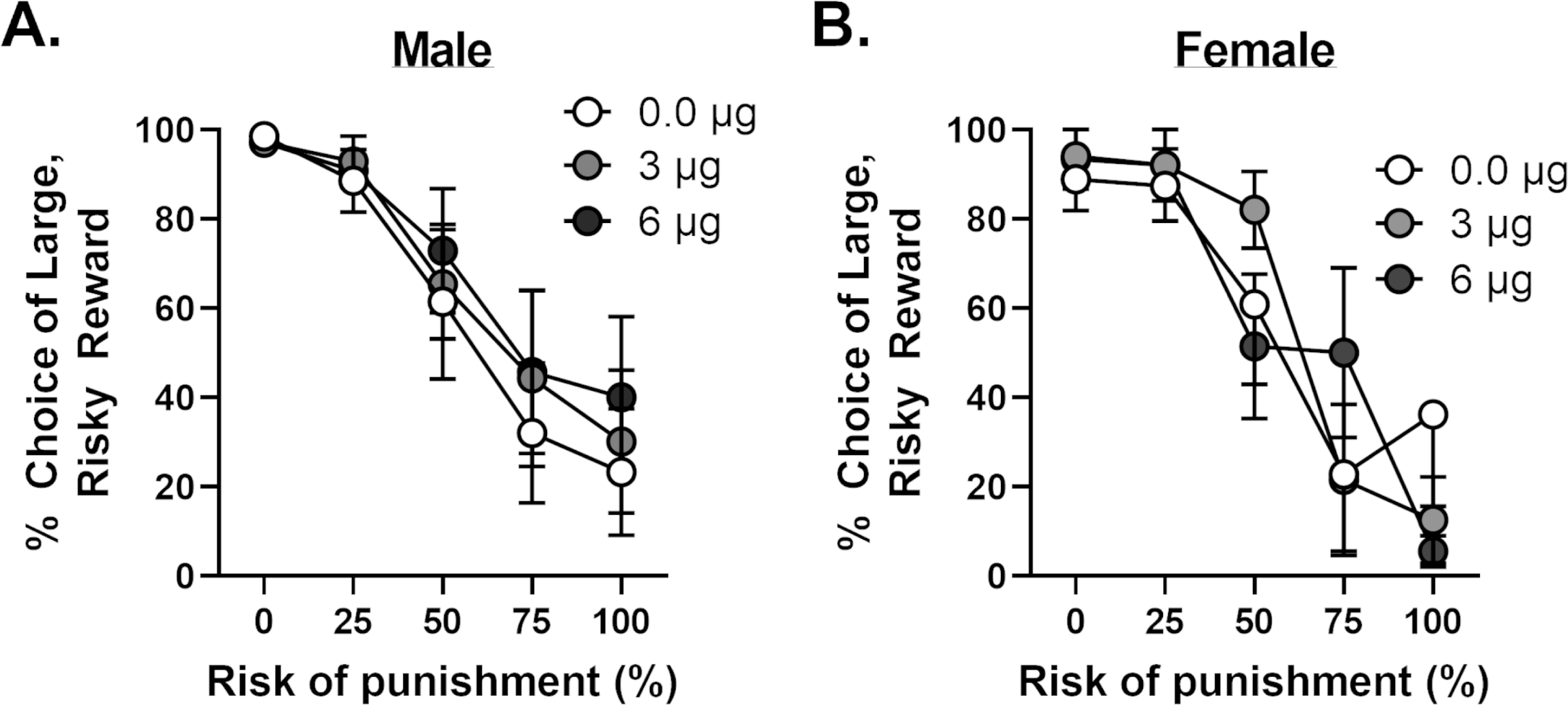
Effects of intra-BLA infusions of a D3R agonist on risk taking. **A.** In males, intra-BLA infusions of the D3R agonist PD128907 did not alter choice of the large, risky reward relative to vehicle infusions. Absence of error bars (e.g., Block 1) is because the size of the standard error of the mean (SEM) was smaller than the point displayed. **B.** In females, intra-BLA infusions of PD128907 did not alter choice of the large, risky reward relative to vehicle infusions. Data are presented as mean ± SEM.

### Experiment 3: The role of BLA D1Rs in risk taking

#### Experiment 3.1

Intra-BLA infusions of the D1R agonist SKF 81297 did not alter choice of the large, risky reward [dose, *F*(2,28)=0.74, *p*=0.49, ƞ^2^=0.05; dose X trial block, *F*(8,112)=0.94, *p*=0.49, ƞ^2^=0.06] in males or females [sex, *F*(1,14)=1.31, *p*=0.27, ƞ^2^=0.07; dose X sex, *F*(2,28)=2.04, *p*=0.15, ƞ^2^=0.13; dose X sex X trial block, *F*(8,112)=1.22, *p*=0.30, ƞ^2^=0.08; Figure 5A, B].

**Figure 5.**
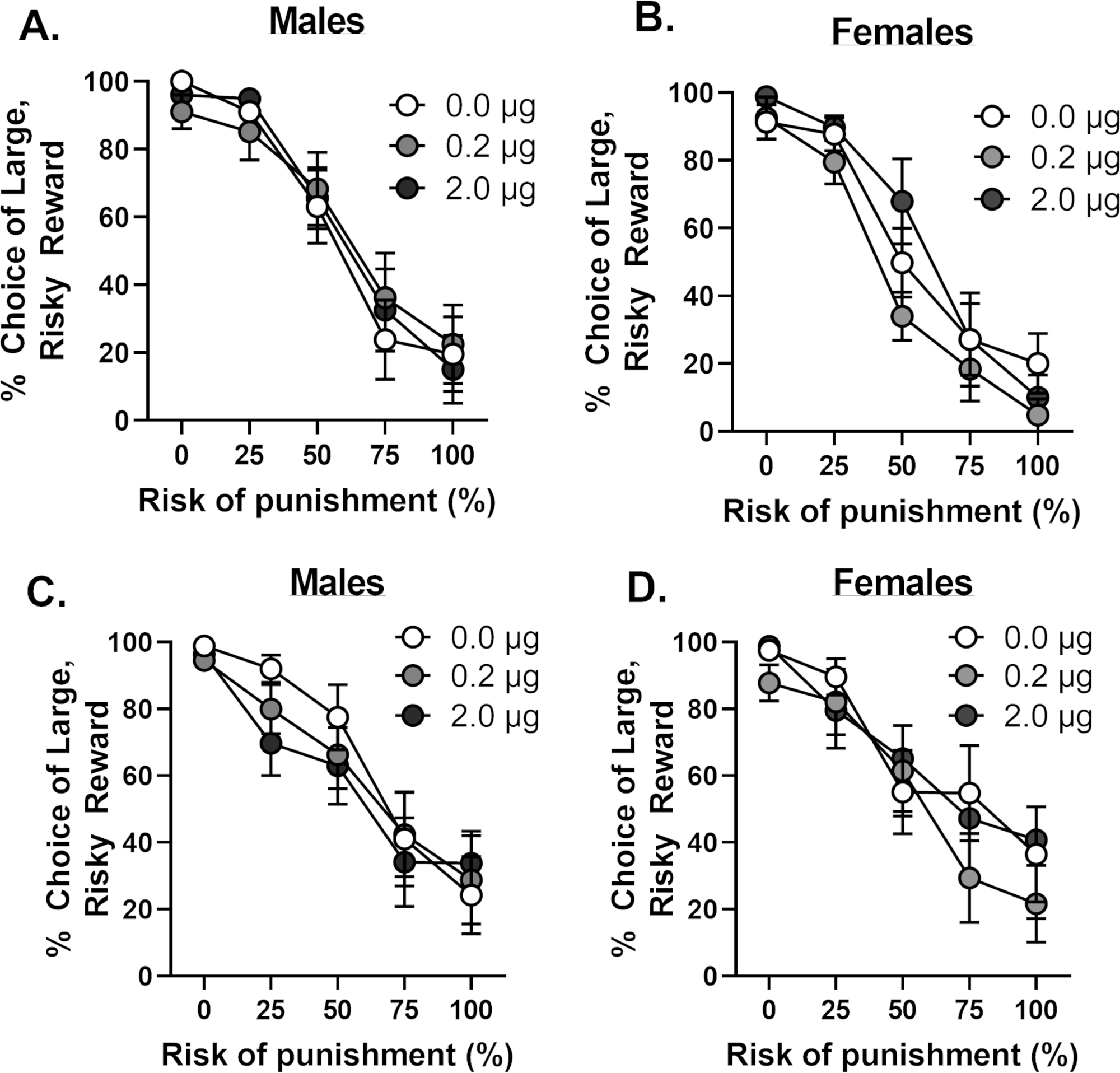
Effects of intra-BLA infusions of D1R ligands on risk taking. **A.** In males, intra-BLA infusions of the D1R agonist SKF81297 did not alter choice of the large, risky reward relative to vehicle infusions. **B.** In females, intra-BLA infusions of SKF81297 did not alter choice of the large, risky reward relative to vehicle infusions. **C.** In males, intra-BLA infusions of the D1R antagonist SCH23390 did not alter choice of the large, risky reward relative to vehicle infusions. **D.** In females, intra-BLA infusions of SCH23390 did not alter choice of the large, risky reward relative to vehicle infusions. Data are presented as mean ± standard error of the mean (SEM). Absence of error bars (e.g., Block 1) is because the size of the SEM was smaller than the point displayed.

#### Experiment 3.2

Similar to the effects of the D1R agonist, intra-BLA infusions of the D1R antagonist SCH23390 did not affect choice of the large, risky reward [dose, *F*(2,24)=0.60, *p*=0.56, ƞ^2^=0.05; dose X trial block, *F*(8,96)=1.49, *p*=0.17, ƞ^2^=0.11] in males or females [sex, *F*(1,12)=0.03, *p*=0.87, ƞ^2^<0.01; dose X sex, *F*(2,24)=0.20, *p*=0.82, ƞ^2^=0.02; dose X sex X trial block, *F*(8,96)=1.56, *p*=0.15, ƞ^2^ = 0.12; Figure 5C, D].

## Discussion

The BLA is a critical node in a network of brain regions involved in decision making involving risk of explicit punishment [30,36–38]. The present findings extend our understanding of the role of BLA in risk taking by revealing the contribution of the dopaminergic (DA) system in the BLA’s ability to guide choice behavior. Consistent with systemic DA manipulations of risk taking [12,13], only D2R agonists reduced risk taking. Surprisingly, however, these effects were sex dependent. Risk taking was reduced in females at doses that were ineffective in altering performance in males. Only when the dose of quinpirole was further increased did risk taking decrease in males. These findings suggest that D2Rs in the BLA of females are more sensitive to DA relative to males, and that this greater sensitivity may contribute to the well-established sex differences in risk taking.

### A role for D2Rs in the BLA in mediating sex differences in risk taking

Prior work has repeatedly demonstrated that females are more risk averse than males [14,39–42], and that this sex difference is largely mediated by a greater sensitivity to risk of punishment in females [42] and dependent on estradiol (E2) [43]. The results of the current study advance our understanding of sex differences in risk taking and reveal potential neural mechanisms that may mediate phenotypical female risk aversion. Our primary finding was that females were more sensitive to intra-BLA infusions of a D2R agonist compared with males. Quinpirole decreased risk taking in females at all doses but was only effective in males at the highest dose. This increased risk aversion in females was driven by an increased propensity to shift to the safe option after receiving a punished large reward (i.e., increased lose-shift; Figure 1C), indicating that D2R activation in the BLA enhanced females’ sensitivity to punishment (note that this effect was not observed in Experiment 1.2, likely due to exclusive preference of the safe reward). Quinpirole also appeared to affect females’ assessment of the risk associated with the large lever during the choice period of the task, as quinpirole-induced risk aversion was accompanied by a greater frequency of abortive large lever presses. Notably, quinpirole also increased abortive lever presses in males at the highest dose. Collectively, these data reveal that activation of D2Rs in the BLA not only provides important negative feedback information to guide subsequent choice toward safer options but is also involved in the initial risk assessment when deciding between available options.

Notably, quinpirole is not selective for D2Rs, but also has binding affinity for D3Rs [44,45], which are also present in the BLA [46]. Hence, the effects of quinpirole on risk taking could be attributable to actions at D3Rs or at D2R and D3Rs in combination. To address this possibility, a selective D3R agonist was infused into the BLA. Activation of D3Rs had no effect on risk taking in either sex, confirming that effects of quinpirole were D2R dependent. These findings are not entirely surprising as systemic administration of the same D3R agonist is ineffective in modulating risk taking in the RDT (although it moderately impairs reward discrimination) [13]. Interestingly, performance in this task is also insensitive to systemic administration of D2R antagonists [12]. This is consistent with the current finding that intra-BLA infusions of a D2R antagonist did not affect risk taking. Given the robust effects of intra-BLA infusions of quinpirole, however, the complete absence of intra-BLA infusions of a D2R antagonist effects is somewhat puzzling. Such disparate effects may be due to the fact that D2Rs are located both presynaptically, serving as autoreceptors, as well as postsynaptically [47]. The consequences of agonists vs. antagonists binding to D2Rs at both sites might differ in terms of their impact on postsynaptic activity and behavioral output. For example, if D2R antagonists bind to D2Rs at DA synapses in the BLA originating from the ventral tegmental area, they would increase DA release by blocking autoreceptors, but simultaneously block postsynaptic receptors, counteracting any potential effect of augmented DA release in the synapse. In contrast, although D2R agonists would decrease DA release by binding to autoreceptors, they would still be able to bind to and stimulate postsynaptic receptors. The pharmacological tools employed in the current study are unable to distinguish between action of ligands at pre- and postsynaptic D2Rs, but these tools used in combination with those that allow real-time measurements of DA release *in vivo* may provide a way to resolve how D2Rs in the BLA regulate risk taking.

Although the sex-dependent effects of quinpirole were initially surprising, they are consistent with recent work showing D2R mRNA in the BLA is higher in females relative to males [48]. Accordingly, augmented sensitivity to risk of punishment in females may be due to either a greater number or greater sensitivity of D2Rs to DA in the BLA. Enhanced D2R sensitivity to DA in the BLA of females may arise from interactions between E2 and DA modulation of BLA activity. Within the BLA, DA can modulate excitability of pyramidal neurons (PNs) by binding to D2Rs located on parvalbumin-expressing interneurons (PV-INs) [20,21,49]. These interneurons directly inhibit BLA PNs; however, the application of DA suppresses the release of GABA from PV-INs onto PNs by binding to D2Rs, resulting in greater excitability of PNs [20,21]. Intriguingly, PV expression varies across the estrous cycle of a rat, with reduced expression of PV in interneurons during phases of the cycle in which E2 levels are high [50]. Because PV expression is activity-dependent [51–53], activity of PV-INS is likely attenuated when there are higher levels of circulating E2. Hence, when E2 levels are elevated, E2 may enhance the sensitivity of D2Rs on PV-INs to DA, possibly via estrogen receptors located on the same interneurons [54], resulting in an inhibition of GABA release from PV-INs. Suppression of PV-INs would lead to a subsequent increase in activity in BLA PNs. This hypothetical model is consistent with studies showing that BLA excitatory activity is not only heightened in females relative to males [50,55], but it is also greater during periods in which E2 levels are elevated [50,56–59]. Consequently, greater risk aversion in females relative to males may be dependent on interactions between E2 and D2Rs in the BLA that result in heightened excitability in the BLA. It is important to note that estrous cycle was not tracked in the current study, and we were therefore unable to determine whether quinpirole was more or less effective in decreasing risk taking depending on the estrous phase. This experiment, as well as those exploring the interaction between D2Rs and E2 in regulating BLA PV-INs, are important next steps to further elucidate the mechanisms underlying sex differences in risk taking.

### Activity of D1Rs in the BLA does not modulate certain forms of risk taking

In contrast to prior studies in which pharmacological manipulations of BLA D1Rs affected performance in a risk-based decision-making task [29], neither a D1R agonist nor a D1R antagonist in the BLA altered risk taking in the RDT. Using a probability discounting task in which rats chose between a small, guaranteed reward and a large reward delivered with varying probabilities, Larkin et al. (2016) showed that intra-BLA infusions of a D1R antagonist decreased choice of the large reward as the probability of reward omission increased. Effects of BLA D1R agonists, however, were more nuanced, as they depended on baseline levels of risk preference, with rats that preferentially chose the large reward being more sensitive to D1R activation. The incongruency between these findings and those of the current study are likely due to the nature of the cost involved in the decision-making task. Indeed, there is a precedent for contrasting effects of pharmacological manipulations on risk-based decision making depending on whether the risk involves punishment (e.g., RDT) versus reward omission (probability discounting task) [30,31]. Notably, the distinct nature of the costs involved in each task may also explain why a D2R agonist decreased risk taking in males in a probability discounting task but had no effect in males in the RDT using identical doses. The lack of BLA D1R involvement in decision making under risk of punishment is consistent, however, with previous behavioral pharmacology studies using the RDT, which showed that performance in the task was insensitive to systemic D1R manipulations [12]. Collectively, the data in the current study add to the growing evidence that decision making under risk of punishment is heavily dependent on D2R activation (although the extent of their involvement appears to differ between males and females) with minimal involvement of D1Rs.

### Conclusions and Future Directions

In summary, the findings in the current study reveal that D2Rs, but not D1Rs, in the BLA have an important role in decision making under risk of punishment. Unexpectedly, female risk taking was more sensitive to D2R activation than males, suggesting differential D2R sensitivity to DA may contribute to sex differences in risk taking. Future studies will focus on identifying whether E2 interacts with D2Rs in the BLA to promote phenotypical female risk aversion. Because activity in the BLA is differentially recruited during distinct aspects of decision making, additional experiments will probe the temporal dynamics of BLA neurons that express D2Rs during risk taking. Collectively, this line of work will provide insight into the neural mechanisms underlying sex differences in risk taking and provide a framework from which to understand psychiatric disorders characterized by pathological decision-making behavior.

## Author contributions

AW designed the work, conducted the experiments, analyzed the data, and drafted the manuscript. LMT and AB conducted the experiments. CAO conceptualized the experiments, aided in their design, analyzed the data, revised the manuscript, and approved the final version. CAO is accountable for all aspects of the work presented in the manuscript.

## Acknowledgements

This research was generously supported by R00DA41493 (CAO), R01DA55676 (CAO) and T32DA018926 (AW) from the National Institute on Drug Abuse.

The authors have nothing to disclose.

## References

1 Brand M, Grabenhorst F, Starcke K, Vandekerckhove MM, Markowitsch HJ. Role of the amygdala in decisions under ambiguity and decisions under risk: evidence from patients with Urbach-Wiethe disease. Neuropsychologia. 2007;45(6):1305–17.

2 Bechara A, Damasio H, Damasio AR, Lee GP. Different contributions of the human amygdala and ventromedial prefrontal cortex to decision-making. The Journal of neuroscience: the official journal of the Society for Neuroscience. 1999;19(13):5473–81.

3 Kahn I, Yeshurun Y, Rotshtein P, Fried I, Ben-Bashat D, Hendler T. The role of the amygdala in signaling prospective outcome of choice. Neuron. 2002;33(6):983–94.

4 Jung WH, Lee S, Lerman C, Kable JW. Amygdala Functional and Structural Connectivity Predicts Individual Risk Tolerance. Neuron. 2018;98(2):394–404 e4.

5 Ferland JN, Adams WK, Murch S, Wei L, Clark L, Winstanley CA. Investigating the influence of ‘losses disguised as wins’ on decision making and motivation in rats. Behavioural pharmacology. 2018;29(8):732–44.

6 Tremblay M, Cocker PJ, Hosking JG, Zeeb FD, Rogers RD, Winstanley CA. Dissociable effects of basolateral amygdala lesions on decision making biases in rats when loss or gain is emphasized. Cogn Affect Behav Neurosci. 2014;14(4):1184–95.

7 Zeeb FD, Winstanley CA. Lesions of the basolateral amygdala and orbitofrontal cortex differentially affect acquisition and performance of a rodent gambling task. The Journal of neuroscience: the official journal of the Society for Neuroscience. 2011;31(6):2197–204.

8 Orsini CA, Trotta RT, Bizon JL, Setlow B. Dissociable Roles for the Basolateral Amygdala and Orbitofrontal Cortex in Decision-Making under Risk of Punishment. The Journal of neuroscience: the official journal of the Society for Neuroscience. 2015;35(4):1368–79.

9 Ghods-Sharifi S, St Onge JR, Floresco SB. Fundamental contribution by the basolateral amygdala to different forms of decision making. The Journal of neuroscience: the official journal of the Society for Neuroscience. 2009;29(16):5251–9.

10 van Holstein M, MacLeod PE, Floresco SB. Basolateral amygdala - nucleus accumbens circuitry regulates optimal cue-guided risk/reward decision making. Progress in neuro-psychopharmacology & biological psychiatry. 2020;98:109830.

11 Orsini CA, Hernandez CM, Singhal S, Kelly KB, Frazier CJ, Bizon JL, et al. Optogenetic Inhibition Reveals Distinct Roles for Basolateral Amygdala Activity at Discrete Time Points during Risky Decision Making. The Journal of neuroscience: the official journal of the Society for Neuroscience. 2017;37(48):11537–48.

12 Simon NW, Montgomery KS, Beas BS, Mitchell MR, LaSarge CL, Mendez IA, et al. Dopaminergic modulation of risky decision-making. The Journal of neuroscience: the official journal of the Society for Neuroscience. 2011;31(48):17460–70.

13 Blaes SL, Orsini CA, Mitchell MR, Spurrell MS, Betzhold SM, Vera K, et al. Monoaminergic modulation of decision-making under risk of punishment in a rat model. Behavioural pharmacology. 2018;29(8):745–61.

14 Orsini CA, Willis ML, Gilbert RJ, Bizon JL, Setlow B. Sex differences in a rat model of risky decision making. Behavioral neuroscience. 2016;130(1):50–61.

15 Simon NW, Gilbert RJ, Mayse JD, Bizon JL, Setlow B. Balancing risk and reward: a rat model of risky decision making. Neuropsychopharmacology: official publication of the American College of Neuropsychopharmacology. 2009;34(10):2208–17.

16 Mitchell MR, Weiss VG, Beas BS, Morgan D, Bizon JL, Setlow B. Adolescent risk taking, cocaine self-administration, and striatal dopamine signaling. Neuropsychopharmacology: official publication of the American College of Neuropsychopharmacology. 2014;39(4):955–62.

17 Asan E. The catecholaminergic innervation of the rat amygdala. Adv Anat Embryol Cell Biol. 1998;142:1–118.

18 Brinley-Reed M, McDonald AJ. Evidence that dopaminergic axons provide a dense innervation of specific neuronal subpopulations in the rat basolateral amygdala. Brain research. 1999;850(1-2):127–35.

19 Asan E. Ultrastructural features of tyrosine-hydroxylase-immunoreactive afferents and their targets in the rat amygdala. Cell Tissue Res. 1997;288(3):449–69.

20 Chu HY, Ito W, Li J, Morozov A. Target-specific suppression of GABA release from parvalbumin interneurons in the basolateral amygdala by dopamine. The Journal of neuroscience: the official journal of the Society for Neuroscience. 2012;32(42):14815–20.

21 Bissiere S, Humeau Y, Luthi A. Dopamine gates LTP induction in lateral amygdala by suppressing feedforward inhibition. Nature neuroscience. 2003;6(6):587–92.

22 Inglis FM, Moghaddam B. Dopaminergic innervation of the amygdala is highly responsive to stress. Journal of neurochemistry. 1999;72(3):1088–94.

23 Harmer CJ, Phillips GD. Enhanced dopamine efflux in the amygdala by a predictive, but not a non-predictive, stimulus: facilitation by prior repeated D-amphetamine. Neuroscience. 1999;90(1):119–30.

24 Hori K, Tanaka J, Nomura M. Effects of discrimination learning on the rat amygdala dopamine release: a microdialysis study. Brain research. 1993;621(2):296–300.

25 Young AM, Rees KR. Dopamine release in the amygdaloid complex of the rat, studied by brain microdialysis. Neuroscience letters. 1998;249(1):49–52.

26 de Oliveira AR, Reimer AE, de Macedo CE, de Carvalho MC, Silva MA, Brandao ML. Conditioned fear is modulated by D2 receptor pathway connecting the ventral tegmental area and basolateral amygdala. Neurobiology of learning and memory. 2011;95(1):37–45.

27 Lintas A, Chi N, Lauzon NM, Bishop SF, Gholizadeh S, Sun N, et al. Identification of a dopamine receptor-mediated opiate reward memory switch in the basolateral amygdala-nucleus accumbens circuit. The Journal of neuroscience: the official journal of the Society for Neuroscience. 2011;31(31):11172–83.

28 Antunes GF, Gouveia FV, Rezende FS, Seno MDJ, de Carvalho MC, de Oliveira CC, et al. Dopamine modulates individual differences in avoidance behavior: A pharmacological, immunohistochemical, neurochemical and volumetric investigation. Neurobiol Stress. 2020;12:100219.

29 Larkin JD, Jenni NL, Floresco SB. Modulation of risk/reward decision making by dopaminergic transmission within the basolateral amygdala. Psychopharmacology. 2016;233(1):121–36.

30 Orsini CA, Moorman DE, Young JW, Setlow B, Floresco SB. Neural mechanisms regulating different forms of risk-related decision-making: Insights from animal models. Neuroscience and biobehavioral reviews. 2015.

31 Winstanley CA, Floresco SB. Deciphering Decision Making: Variation in Animal Models of Effort- and Uncertainty-Based Choice Reveals Distinct Neural Circuitries Underlying Core Cognitive Processes. The Journal of neuroscience: the official journal of the Society for Neuroscience. 2016;36(48):12069–79.

32 Farrell MR, Ruiz CM, Castillo E, Faget L, Khanbijian C, Liu S, et al. Ventral pallidum is essential for cocaine relapse after voluntary abstinence in rats. Neuropsychopharmacology: official publication of the American College of Neuropsychopharmacology. 2019;44(13):2174–85.

33 Halladay LR, Kocharian A, Piantadosi PT, Authement ME, Lieberman AG, Spitz NA, et al. Prefrontal Regulation of Punished Ethanol Self-administration. Biological psychiatry. 2020;87(11):967–78.

34 Blanchard DC, Griebel G, Pobbe R, Blanchard RJ. Risk assessment as an evolved threat detection and analysis process. Neuroscience and biobehavioral reviews. 2011;35(4):991–8.

35 Hunt HF, Brady JV. Some effects of punishment and intercurrent anxiety on a simple operant. J Comp Physiol Psychol. 1955;48(4):305–10.

36 Piantadosi PT, Halladay LR, Radke AK, Holmes A. Advances in understanding meso-cortico-limbic-striatal systems mediating risky reward seeking. Journal of neurochemistry. 2021;157(5):1547–71.

37 Wassum KM, Izquierdo A. The basolateral amygdala in reward learning and addiction. Neuroscience and biobehavioral reviews. 2015;57:271–83.

38 Liley AE, Gabriel DBK, Simon NW. Lateral Orbitofrontal Cortex and Basolateral Amygdala Regulate Sensitivity to Delayed Punishment during Decision-making. eNeuro. 2022;9(5).

39 Liley AE, Gabriel DBK, Sable HJ, Simon NW. Sex Differences and Effects of Predictive Cues on Delayed Punishment Discounting. eNeuro. 2019;6(4).

40 Chowdhury TG, Wallin-Miller KG, Rear AA, Park J, Diaz V, Simon NW, et al. Sex differences in reward- and punishment-guided actions. Cogn Affect Behav Neurosci. 2019;19(6):1404–17.

41 Islas-Preciado D, Wainwright SR, Sniegocki J, Lieblich SE, Yagi S, Floresco SB, et al. Risk-based decision making in rats: Modulation by sex and amphetamine. Hormones and behavior. 2020;125:104815.

42 Truckenbrod LM, Cooper EM, Orsini CA. Cognitive mechanisms underlying decision making involving risk of explicit punishment in male and female rats. Cogn Affect Behav Neurosci. 2022.

43 Orsini CA, Blaes SL, Hernandez CM, Betzhold SM, Perera H, Wheeler AR, et al. Regulation of risky decision making by gonadal hormones in males and females. Neuropsychopharmacology: official publication of the American College of Neuropsychopharmacology. 2021;46(3):603–13.

44 Levant B, Grigoriadis DE, DeSouza EB. [3H]quinpirole binding to putative D2 and D3 dopamine receptors in rat brain and pituitary gland: a quantitative autoradiographic study. The Journal of pharmacology and experimental therapeutics. 1993;264(2):991–1001.

45 Malmberg A, Mohell N. Characterization of [3H]quinpirole binding to human dopamine D2A and D3 receptors: effects of ions and guanine nucleotides. The Journal of pharmacology and experimental therapeutics. 1995;274(2):790–7.

46 Bouthenet ML, Souil E, Martres MP, Sokoloff P, Giros B, Schwartz JC. Localization of dopamine D3 receptor mRNA in the rat brain using in situ hybridization histochemistry: comparison with dopamine D2 receptor mRNA. Brain research. 1991;564(2):203–19.

47 Beaulieu JM, Gainetdinov RR. The physiology, signaling, and pharmacology of dopamine receptors. Pharmacological reviews. 2011;63(1):182–217.

48 Georgiou P, Zanos P, Bhat S, Tracy JK, Merchenthaler IJ, McCarthy MM, et al. Dopamine and Stress System Modulation of Sex Differences in Decision Making. Neuropsychopharmacology: official publication of the American College of Neuropsychopharmacology. 2018;43(2):313–24.

49 Pinard CR, Muller JF, Mascagni F, McDonald AJ. Dopaminergic innervation of interneurons in the rat basolateral amygdala. Neuroscience. 2008;157(4):850–63.

50 Blume SR, Freedberg M, Vantrease JE, Chan R, Padival M, Record MJ, et al. Sex- and Estrus-Dependent Differences in Rat Basolateral Amygdala. The Journal of neuroscience: the official journal of the Society for Neuroscience. 2017;37(44):10567–86.

51 Philpot BD, Lim JH, Brunjes PC. Activity-dependent regulation of calcium-binding proteins in the developing rat olfactory bulb. J Comp Neurol. 1997;387(1):12–26.

52 Jiao Y, Zhang C, Yanagawa Y, Sun QQ. Major effects of sensory experiences on the neocortical inhibitory circuits. The Journal of neuroscience: the official journal of the Society for Neuroscience. 2006;26(34):8691–701.

53 Carder RK, Leclerc SS, Hendry SH. Regulation of calcium-binding protein immunoreactivity in GABA neurons of macaque primary visual cortex. Cerebral cortex. 1996;6(2):271–87.

54 Blurton-Jones M, Tuszynski MH. Estrogen receptor-beta colocalizes extensively with parvalbumin-labeled inhibitory neurons in the cortex, amygdala, basal forebrain, and hippocampal formation of intact and ovariectomized adult rats. J Comp Neurol. 2002;452(3):276–87.

55 Lebron-Milad K, Abbs B, Milad MR, Linnman C, Rougemount-Bucking A, Zeidan MA, et al. Sex differences in the neurobiology of fear conditioning and extinction: a preliminary fMRI study of shared sex differences with stress-arousal circuitry. Biol Mood Anxiety Disord. 2012;2:7.

56 Blume SR, Padival M, Urban JH, Rosenkranz JA. Disruptive effects of repeated stress on basolateral amygdala neurons and fear behavior across the estrous cycle in rats. Sci Rep. 2019;9(1):12292.

57 Zeidan MA, Igoe SA, Linnman C, Vitalo A, Levine JB, Klibanski A, et al. Estradiol modulates medial prefrontal cortex and amygdala activity during fear extinction in women and female rats. Biological psychiatry. 2011;70(10):920–7.

58 Macoveanu J, Henningsson S, Pinborg A, Jensen P, Knudsen GM, Frokjaer VG, et al. Sex-Steroid Hormone Manipulation Reduces Brain Response to Reward. Neuropsychopharmacology: official publication of the American College of Neuropsychopharmacology. 2016;41(4):1057–65.

59 van Wingen GA, Ossewaarde L, Backstrom T, Hermans EJ, Fernandez G. Gonadal hormone regulation of the emotion circuitry in humans. Neuroscience. 2011;191:38–45.

