## Supplemental Materials for "Sex differences in sensitivity to dopamine receptor manipulations of risk-based decision making in rats"

**METHODS**

*Subjects*

Male and female Long-Evans rats were obtained from Charles River Laboratories (Hollister, CA), arriving on postnatal day 60. Rats were housed in ventilated cages with Sani-Chip bedding and provided Nyla bones and nestlets for enrichment. Prior to experimental procedures, rats were handled once a day for a minimum of 3 days to habituate the rats to the experimenters. Rats were food restricted during behavioral training, with target weights adjusted upward by 5 g each week to account for growth. Once fully grown (~250 g for females and ~350 g for males), rats were fed 8 g of soy-free food (Envigo Teklad Irradiated Global 19% Protein Extruded Rodent Diet, #2919) per day.

*Apparatus*

During behavioral sessions, rats were tested in six identical standard operant chambers (Coulbourn Instruments, Whitehall, PA), each of which were housed within a single sound-attenuating cabinet (Coulbourn Instruments) outfitted with red lights and noise insulation foam. In each operant chamber, a food trough was positioned in the middle of the front wall, extending 3 cm into the chamber and containing a photobeam to detect entries into the food trough. Food pellets (45 mg; Lab Supply, 5UTL, Houston, TX) were delivered into the food trough via its connection to a feeder. A nosepoke hole was located directly above the food trough and was flanked on either side by two retractable levers. A 1.12W light was mounted on the back of the cabinet and was used as a house light during behavioral testing. The floor of the test chambers consisted of stainless-steel rods connected to a shock generator (Coulbourn Instruments) that delivered scrambled footshocks to the floor. Operant chambers were interfaced with a computer running Graphic State 4 software, which concurrently controlled chamber components, such as levers and food delivery, and recorded task events for data analysis. Chambers were cleaned with diluted Nolvasan disinfectant between testing of successive groups of rats.

*Surgical procedures*

One week after arrival, rats underwent stereotaxic surgery in which bilateral cannulae were implanted directly above the basolateral amygdala (BLA). Rats were anesthetized with isoflurane gas (1-5% in O_2_) and were administered Meloxicam (2mg/kg), buprenorphine (0.03mg/kg), and sterile saline (10mL) subcutaneously before being placed in the stereotaxic apparatus (David Kopf). After placing an adhesive surgical drape on the rat’s body, the scalp was disinfected with a chlorohexidine/isopropyl alcohol wipe and 2% lidocaine was injected subcutaneously under the skin. The scalp was incised and retracted to reveal the rat’s skull, and four small burr holes were drilled into the skull for jeweler’s screws. Screws were placed such that two would be anterior and two would be posterior to the guide cannulae. The skull was then leveled to ensure that bregma and lambda were in the same horizontal plane. Two additional burr holes were drilled for bilateral implantation of guide cannulae (Plastics One; 22 gauge) above the BLA (AP: -3.3, ML: ±4.9, DV: -7.3 from skull surface). Cannulae were anchored into place with Metabond and dental cement and the incision was sutured closed after the dental cement hardened. Sterile stylets were inserted into the cannulae to prevent occlusion during recovery and behavioral training. Following surgery, rats were placed back in their homecage on a heating pad to recover and were administered an additional 10 mL of sterile saline subcutaneously. Rats recovered for one week, after which they were food restricted for behavioral testing. Stylets were replaced with new sterile stylets on a weekly basis until microinfusions began.

*Behavioral procedures*

Shaping

Rats were first trained to perform individual components of the Risky Decision-making Task (RDT), such as nosepoking and lever pressing for food delivery. In the initial shaping session, rats learned to retrieve a single food pellet, delivered every 100 ± 40 seconds, from the food trough. Upon meeting the criteria of 100 food trough entries in the 64-minute session, rats progressed to lever shaping protocols. In these sessions, a single lever (left or right, counterbalanced across rats) was extended into the chamber and remained so for the entire 30-minute session. Rats learned that a lever press on the extended lever resulted in the delivery of a single food pellet. After reaching passing criterion on one lever (50 lever presses in the 30-minute session), rats were shaped to lever press on the opposite lever and were required to reach the same passing criterion used for the first lever shaping session. After successful lever shaping, rats were trained to nosepoke in the nosepoke hole positioned directly above the food trough. A nosepoke triggered the extension of one of the two levers (randomly determined), a press on which resulted in lever retraction, the termination of the house light and delivery of a single food pellet. To progress to the final phase of shaping, rats were required to lever press 30 times on each lever in the 60-minute session.

Reward Discrimination

Prior to training on the RDT, rats learned to discriminate between a small food reward (1 pellet) and a large food reward (2 pellets) in a Reward Discrimination (RD) task. Each 40-second trial began with the illumination of the houselight and nosepoke hole. A poke into the nosepoke hole extinguished the light in the nosepoke hole and resulted in the extension of a single lever (i.e., forced choice trials) or both levers (i.e., free choice trials) into the chamber. If a rat failed to nosepoke within 10 seconds, the house light and nosepoke lights were extinguished and the trial was scored as an omission. A press on one lever yielded the delivery of 1 food pellet whereas a press on the other lever yielded the delivery of 2 food pellets. The identity of the lever (small vs. large) was counterbalanced across sexes and remained consistent throughout training on both RD and the RDT. If a rat failed to lever press within 10 seconds, the house light was extinguished, the lever(s) were retracted, and the trial was scored as an omission. If a rat successfully lever pressed, lever(s) were retracted and the house light was extinguished for the remainder of the trial. This task was 60 minutes in duration and consisted of 5 blocks of 18 trials. Each block began with 8 forced choice trials in which a single lever was extended (4 trials per lever; randomized across the 8 trials) and was followed by 10 free choice trials in which both levers were extended, and rats could choose freely between them. Rats were trained on this task until they demonstrated preference for the large reward (≥80% choice of the large lever) for 3 consecutive days.

Risky Decision-making Task

Upon meeting passing criteria on RD, rats began training on the RDT until they reached stable choice performance (see Data Analysis for a definition of stable behavior). The RDT was similar in structure to RD with the exception that the delivery of the large reward was associated with the probability of the delivery of a 1-second footshock. The probability of footshock systematically increased across the five blocks of trials in 25% increments (0, 25, 50, 75, 100%). The forced choice trials served to inform the rat about the risk contingencies in effect for that trial block. The probability of footshock on each forced choice trial was dependent across the four forced choice trials associated with the large, risky lever. For example, on the 25% trial block, only one of the four lever presses on the large, risky lever would result in the delivery of a footshock. In contrast, on the 75% trial block, three out of the four lever presses on the large, risky lever would result in the delivery of a footshock. Unlike forced choice trials, the probability of footshock on the free choice trials was independent of the outcomes of other free choice trials in that block (i.e., the probability of footshock delivery was equivalent across the free choice trials in the block). Regardless of footshock probability, two food pellets were delivered every time the rat pressed the large, risky lever. Shock intensities were initially set at 0.20 mA for males and 0.15 mA for females but were then adjusted individually for each rat throughout the duration of training. These adjustments were necessary to ensure that mean baseline performance was close to the center of the parametric space and thus improve the likelihood of observing increases or decreases in risk taking as a result of pharmacological manipulations. Once rats reached behavioral stability, rats received intra-BLA microinfusions of dopamine receptor targeting drugs.

*Microinfusions*

Upon reaching behavioral stability, rats underwent mock microinjection procedures in which injectors (Plastics One; 28 gauge) were inserted into the guide cannulae, but no drug was microinfused. Rats were placed back in their homecage for 10 minutes and then tested on the RDT. This procedure was conducted to ensure that the insertion of the injector alone (and any associated mechanical damage) had no effect on choice behavior. Forty-eight hours later, rats received intra-BLA microinfusions (0.5 µl at a rate of 0.4 µl/min in each hemisphere) of quinpirole (Experiments 1.1 and 1.2), eticlopride (Experiment 1.3), PD 128907 (Experiment 2), SKF 81297 (Experiment 3.1), or SCH 23390 (Experiment 3.2). In each experiment, the different doses of each drug were microinfused based on a randomized, within-subjects design such that each rat received each dose of the drug and its corresponding vehicle. Each successive infusion of a drug dose was separated by minimum of 48 hours. Following each microinfusion, the injectors remained in place for an additional minute to allow for drug diffusion, after which injectors were removed from the guide cannulae and replaced with sterile stylets. With the exception of rats in Experiment 1.2, rats were then returned to their homecage for 10 minutes before being tested on the RDT. In Experiment 1.2, the window of time between infusions and testing was extended to 15 minutes due to the significant number of omissions that were observed with intra-BLA infusions of the high dose of quinpirole in initial pilot experiments. The microinjectors extended 1 mm beyond the tip of the guide cannulae and were connected to 10 µl Hamilton syringes via polyethylene (PE-20; Plastics One) tubing. These syringes were mounted on an infusion pump (Harvard Apparatus), which controlled the volume and timing of the infusion.

*Drugs*

In Experiments 1.1 and 1.2, the D2R agonist (-)-quinpirole hydrochloride (0.0, 0.2, 2.0, and 4.0 µg; Tocris Biosciences) was dissolved in 0.9% saline and microinfused into the BLA. In Experiment 1.3, the D2R antagonist eticlopride hydrochloride (0.0, 0.2, and 2.0 µg; Sigma-Aldrich) was dissolved in 0.9% saline and microinfused into the BLA. In Experiment 2, the selective D3R agonist (+)-PD 128907 hydrochloride (0.0, 3.0, and 6.0 µg; Tocris Biosciences) was dissolved in 0.9% saline and microinfused into the BLA. Finally, the D1R agonist SKF 81297 hydrobromide (Experiment 3.1) and D1R antagonist SCH 23390 hydrochloride (Experiment 3.2) were dissolved in 0.9% saline and microinfused into the BLA. Doses for each drug were based on previous studies showing that microinfusions of these drugs at these doses in the BLA or in the nucleus accumbens were effective in altering decision-making behavior in the probability discounting task [1,2]. In all experiments, 0.9% saline served as the vehicle control.

*Histology*

At the end of each experiment, rats were overdosed with Euthasol and perfused intracardially with 0.1M PBS followed by 4% paraformaldehyde. Brains were then extracted and post-fixed in 4% paraformaldehyde for 24 hours and were then transferred to a 20% sucrose in 0.1M PBS solution for a minimum of 2 days before being sectioned. Brains were sectioned at 40 µm on a cryostat maintained at -19°C, and tissue sections were mounted on electrostatic slides (Fisherbrand). Brain sections were then stained with 0.25% thionin and coverslipped with Permount (Fisher Scientific) to verify cannulae placement.

*Data analysis*

Using G*Power Software, power analyses were conducted to determine sample sizes necessary to detect effect sizes of 0.8 or greater, assuming an α of 0.05. Raw data files were extracted and then analyzed using customized Graphic State 4 analysis template for behavioral measures on the RDT. Extracted data were compiled in Microsoft Excel and then exported into SPSS 27 for statistical analyses. GraphPad Prism 9.0 was used to create figures. In all statistical analyses, results were considered significant when *p* ≤ 0.05. If parent ANOVAs yielded significant main effects or interactions, additional post-hoc ANOVAs were conducted to identify the source of the significance, and *p*-values were adjusted to account for multiple comparisons using Bonferroni’s corrections. Effect sizes are reported as ƞ^2^ for ANOVAs and as the absolute value of Cohen’s d for *t*-tests.

The main dependent variables used to analyze performance on the RDT were the percentage of free choice trials in each block on which the rat chose the large, risky lever (risk taking). To determine behavioral stability on the RDT, a three-factor (sex X day X trial block) repeated measures analysis of variance (RMANOVA) was used to analyze risk taking (using both dependent variables) across a sliding window of 3 consecutive behavioral sessions. If these analyses yielded a main effect of trial block (i.e., 0, 25, 50, 75, 100%), but no main effect of day or significant interactions between day, trial block and sex, behavior was considered stable. Effects of pharmacological manipulations on percent choice were evaluated using a RMANOVA, with trial block and drug dose as the within-subjects factors and sex as the between-subjects factor. Similar RMANOVAs were also conducted for each sex separately. If these parent ANOVAs revealed a significant main effect of dose and/or an interaction between dose and trial block, additional RMANOVAs were conducted to compare individual dose(s) with the vehicle treatment. To fully examine the effects of pharmacological manipulations on risk taking, trial-by-trial analyses were conducted to determine whether DA receptor activation or inhibition altered the degree to which feedback from a previous trial affected choice in the subsequent trial. Win-stay behavior, which served as a proxy for sensitivity to rewarding outcomes, was determined by dividing the number of free choice trials on which a rat chose the large, risky lever after receiving the large, unpunished reward (i.e., large reward without footshock) by the total number of free choice trials on which the rat received the large, unpunished reward. Lose-shift behavior, which served as a proxy for sensitivity to punishment, was determined by dividing the number of free choice trials on which a rat chose the small, safe reward after receiving a large, punished reward (i.e., large reward accompanied by footshock) by the total number of free choice trials on which the rat received the large, punished reward. Win-stay and lose-shift behavior were analyzed using a RMANOVA, with dose as the within-subjects factor and sex as the between-subjects factor. These analyses were also conducted for each sex separately.

The effects of DA receptor manipulations on other ancillary behavioral measures on the RDT were also evaluated. Latencies to press levers during forced choice trials (defined as the duration of time between lever extension and a lever press, excluding trial omissions) were analyzed with a four-way RMANOVA, with dose, lever identity (small, safe vs. large, risky) and trial block as the within-subjects factors and sex as the between-subjects factor. These analyses were also conducted for each sex separately. If these parent ANOVAs yielded significant main effects or interactions, additional ANOVAs were conducted to compare individual doses with the vehicle condition for each lever. Finally, trial omissions were defined as the percentage of free choice trials on which a rat failed to lever press in the allotted time. Effects of pharmacological manipulations on trial omissions were assessed using a RMANOVA, with dose as the within-subjects factor and sex as the between-subjects factor. This analysis was also conducted for each sex separately.

**ADDITIONAL RESULTS**

*Experiment 1: The role of D2Rs in the BLA in risk taking*

Histology

Figure S1 depicts the placements of cannulae in the basolateral amygdala for rats included in Experiment 1. In Experiment 1.1, seven out of 30 rats were euthanized prior to beginning microinfusions due to illness. Placements for two rats were too ventral, and these rats were therefore excluded from analysis. Consequently, the final sample size for Experiment 1.1 was n = 21 (n = 11, female; n = 10, male). In Experiment 1.2, five out of 20 rats were euthanized prior to the start of microinfusions due to illness, and an additional three rats were excluded from data analysis due to missed cannulae placements (too ventral). After accounting for attrition and these exclusions, the final sample size for Experiment 1.2 was n = 12 (n = 7, female; n = 6, male). In Experiment 1.3, the same rats in Experiment 1.1 were used; hence, the final sample size for Experiment 1.3 was n = 21 (n = 11, female; n = 10, male).

Behavior:

*Experiment 1.1:*

Although quinpirole did not affect choice behavior in males, latencies to lever press during forced choice trials were still analyzed for each sex separately. In males, there was no effect of quinpirole on latencies to press either lever at any dose relative to vehicle [dose, *F*(2,18)=1.98, *p*=0.17, ƞ^2^=0.18; dose X lever identity, *F*(2,18)=0.44, *p*=0.65, ƞ^2^=0.05]. In females, however, there was a main effect of dose [*F*(2,18)=3.58, *p*=0.05, ƞ^2^=0.28], but no significant interaction between dose and lever identity [*F*(2,18)=1.12, *p*=0.35, ƞ^2^=0.11]. Additional analyses revealed that this main effect of dose was driven by an overall increase in latencies to press levers at the low dose [*F*(1,9)=6.19, *p*=0.04, ƞ^2^=0.41]; there was no effect of the high dose of quinpirole on latencies to press levers [*F*(1,9)=3.04, *p*=0.12, ƞ^2^=0.25].

*Experiment 1.2:*

Because there were no sex differences in the effects of a higher dose of quinpirole on risk taking, latency and omission data were not analyzed separately for each sex. There was no effect of quinpirole on latencies to press either lever relative to vehicle in either sex [dose, *F*(1,10)=3.26, *p*=0.10, ƞ^2^=0.25; sex, *F*(1,10)=0.42, *p*=0.53, ƞ^2^=0.04; dose X lever identity, *F*(1,10)=0.07, *p*=0.80, ƞ^2^<0.01; dose X sex, *F*(1,10)=0.48, *p*=0.51, ƞ^2^=0.05; dose X lever identity X sex, *F*(1,10)=0.36, *p*=0.56, ƞ^2^=0.04].

*Experiment 1.3:*

Eticlopride did not affect latencies to press either lever during forced choice trials in males or females [dose, *F*(2,36)=1.51, *p*=0.24; ƞ^2^=0.08; sex, *F*(1,18)=3.70, *p*=0.07, ƞ^2^=0.17; dose X sex, *F* (2,36)=0.99, *p*=0.38, ƞ^2^=0.05; dose X lever identity, *F*(2,36)=0.36, *p*=0.70, ƞ^2^=0.02; dose X sex X lever identity, *F*(2,36)=1.32, *p*=0.28, ƞ^2^=0.07]. Although females omitted significantly more free choice trials than males [*F*(1,18)=11.44, *p*<0.01, ƞ^2^=0.39], eticlopride did not reliably alter this pattern of behavior [dose, *F*(2,36)=0.98, *p*=0.39, ƞ^2^=0.05; dose X sex, *F*(2,36)=0.53, *p*=0.59, ƞ^2^=0.03].

*Experiment 2: The role of D3Rs in the BLA in risk taking*

Histology

Figure S2 displays the cannulae placements in the basolateral amygdala for rats included in Experiment 2. Thirty-seven rats (twelve of which were also used in Experiment 1.2) underwent surgery, but a significant number (n = 11) became acutely ill during recovery and were quickly euthanized. Two additional rats lost their cranial implant over the course of behavioral training. Out of the remaining 24 rats, five were excluded due to missed placements, resulting in a final sample size of n = 19 (n = 8, female; n = 11, male).

Behavior

Intra-BLA infusions of PD128907 did not alter latencies to press either lever during forced choice trials in either sex [dose, *F*(2,32)=0.44, *p*=0.65, ƞ^2^=0.03; sex, *F*(1,16)<0.01, *p*=0.96, ƞ^2^<0.01; dose X lever identity, *F*(2,32)=0.30, *p*=0.75, ƞ^2^=0.02; dose X sex, *F*(2,32)=2.41, *p*=0.11, ƞ^2^=0.13; dose X sex X lever identity, *F*(2,32)=2.54, *p*=0.09, ƞ^2^=0.14]. Similar to Experiment 1.3, females made significantly more omissions than males [*F*(1,17)=17.46, *p*<0.01, ƞ^2^=0.51], but PD128907 was ineffective in altering this behavior in either sex [dose, *F*(2,34)=0.36, *p*=0.70, ƞ^2^=0.02; dose X sex, *F*(2,34)=0.03, *p*=0.97, ƞ^2^<0.01].

*Experiment 3: The role of D1Rs in the BLA in risk taking*

Histology

Figure S3 depicts the cannulae placements in the basolateral amygdala for rats included in Experiment 3. Rats from Experiment 2 were also used for both Experiments 3.1 and 3.2; eight additional rats underwent surgery for Experiment 3, but three were euthanized during training in the RDT due to illness. The final sample size for Experiment 3.1 and Experiment 3.2 was n = 18 (n = 8, female; n = 10, male).

Behavior

*Experiment 3.1:*

There were no effects of SKF 81297 on latencies to press either lever during forced choice trials in either sex [dose, *F*(2,32)=0.21, *p*=0.81, ƞ^2^=0.01; sex, *F*(1,16)=2.72, *p*=0.12, ƞ^2^=0.15; dose X lever identity, *F*(2,32)=0.24, *p*=0.79, ƞ^2^=0.02; dose X sex, *F*(2,32)=0.17, *p*=0.85, ƞ^2^=0.01; dose X lever identity X sex, *F*(2,32)=0.18, *p*=0.83, ƞ^2^=0.01]. Finally, although females omitted significantly more free choice trials compared with males [*F*(1,16)=8.58, *p*=0.01, ƞ^2^=0.35], SKF 81297 did not alter omissions in either sex [dose, *F*(2,32)=2.09, *p*=0.14, ƞ^2^=0.12; dose X sex, *F*(2,32)=0.16, *p*=0.85, ƞ^2^=0.01].

*Experiment 3.2:*

Although latencies to press levers were greater in females overall relative to males [*F*(1,16)=5.14, *p*=0.04, ƞ^2^=0.24], there was no effect of intra-BLA SCH 23390 on latencies to press levers in the forced choice trials in either sex [dose, *F*(2,32)=0.37, *p*=0.69, ƞ^2^=0.02; dose X lever identity, *F*(2,32)=0.54, *p*=0.59, ƞ^2^=0.03; dose X sex, *F*(2,32)=0.19, *p*=0.83, ƞ^2^=0.01; dose X sex X lever identity, *F*(2,32)=1.61, *p*=0.22, ƞ^2^=0.09]. Consistent with the previous experiments, females made significantly more omissions of free choice trials [*F*(1,16)=16.10, *p*<0.01, ƞ^2^=0.50]. Infusions of SCH 23390, however, did not change the percentage of omissions relative to vehicle infusions in either sex [dose, *F*(2,32)=2.55, *p*=0.09, ƞ^2^=0.14; dose X sex, *F*(2,32)=0.92, *p*=0.41, ƞ^2^=0.05].

**Figure Captions**

***Figure S1. Experiment 1 Histology.* A.** Representation of cannulae placements in the basolateral amygdala in Experiment 1.1. **B.** Representation of cannulae placements in the basolateral amygdala in Experiment 1.2. **C.** Representation of cannulae placements in the basolateral amygdala in Experiment 1.3.

***Figure S2. Experiment 2 Histology.*** Representation of cannulae placements in the basolateral amygdala in Experiment 2.

***Figure S3. Experiment 3 Histology.* A.** Representation of cannulae placements in the basolateral amygdala in Experiment 3.1. **B.** Representation of cannulae placements in the basolateral amygdala in Experiment 3.2.

**References**

1	Larkin JD, Jenni NL, Floresco SB. Modulation of risk/reward decision making by dopaminergic transmission within the basolateral amygdala. Psychopharmacology. 2016;233(1):121-36.

2	Stopper CM, Khayambashi S, Floresco SB. Receptor-specific modulation of risk-based decision making by nucleus accumbens dopamine. Neuropsychopharmacology : official publication of the American College of Neuropsychopharmacology. 2013;38(5):715-28.
